# Rapid, ultra-local adaptation facilitated by phenotypic plasticity

**DOI:** 10.1101/598292

**Authors:** Syuan-Jyun Sun, Andrew M. Catherall, Sonia Pascoal, Benjamin J. M. Jarrett, Sara E. Miller, Michael J. Sheehan, Rebecca M. Kilner

**Affiliations:** Department of Zoology, University of Cambridge, Downing Street, Cambridge, CB2 3EJ, UK; Department of Entomology, Michigan State University, 288 Farm Lane Room 243, East Lansing, MI 48824, USA; Department of Neurobiology and Behavior, Cornell University, 215 Tower Road, Ithaca, NY 14853, USA

**Keywords:** interspecific competition, phenotypic plasticity, plasticity-led evolution, burying beetles, *Nicrophorus vespilloides*, niche expansion, local adaptation

## Abstract

Models of ‘plasticity-first’ evolution are attractive because they explain the rapid evolution of new complex adaptations. Nevertheless, it is unclear whether plasticity can still facilitate rapid evolution when diverging populations are connected by gene flow. Here we show how plasticity has generated adaptive divergence in fecundity in wild populations of burying beetles *Nicrophorus vespilloides*, which are still connected by gene flow, which occupy distinct Cambridgeshire woodlands that are just 2.5km apart and which diverged from a common ancestral population c. 1000-4000 years ago. We show that adaptive divergence is due <u>to</u> the coupling of an evolved increase in the elevation of the reaction norm linking clutch size to carrion size (i.e. genetic accommodation) with plastic secondary elimination of surplus offspring. Working in combination, these two processes have facilitated rapid adaptation to fine-scale environmental differences, despite ongoing gene flow.

## Introduction

Understanding the scale and pace of local adaptation is a long-standing problem in evolutionary biology that has acquired new significance for predicting how natural populations will respond to man-made environmental change^1–6^. In theory, phenotypic plasticity can potentially both accelerate the pace of evolution, and fine-tune the scale of local adaptation, through two distinct processes. The extent of phenotypic plasticity in any trait is described by a reaction norm, which relates environmental variation to the phenotype it induces. ‘Plasticity-first’ evolution can speed up the pace of evolutionary change in complex traits because the shape, slope and elevation of a reaction norm each have genetic components, upon which selection can act^1–6^. Recent experimental work on natural populations has shown that selection in a new environment can greatly reduce the slope of the reaction norm, causing the constitutive expression of novel, canalised traits^7,8^. New adaptations can therefore evolve through the ‘genetic assimilation’ (*sensu*^9^) of traits that were once induced environmentally.

Genetic assimilation yields rapid change but is predicted to occur only when there is little or no gene flow between populations that inhabit different local environments^10–12^. While gene flow still persists, theory predicts that selection will instead act to maintain plasticity in adaptive traits^11,12^. Plasticity is adaptive under these conditions because it ‘forgives’ costly mistakes incurred through the expression of locally maladaptive traits. Nevertheless, local adaptation through genetic change is still possible, if selection fine-tunes the slope or elevation of the reaction norm within each population to match local conditions through genetic accommodation^13,14^. However, whether this mechanism is at work in natural animal populations is unclear (but see^15^ for evidence from a plant species).

We tested whether plasticity has facilitated recent, ultra-local adaptation in wild populations of burying beetles *Nicrophorus vespilloides.* We focused on populations occupying Gamlingay Wood and Waresley Wood in Cambridgeshire, UK, which are c. 2.5 km apart. We have previously shown that the two beetle populations cannot be differentiated at neutral genetic markers, indicating there is ongoing gene flow^16^. Gamlingay Wood and Waresley Wood have existed as distinct woodlands since at least 1086 because they are both recorded in the Domesday Book (a land survey commissioned by William the Conqueror). However, until 3000-4000 years they were almost certainly connected as part of the ‘Wild Wood’, the ancient forest that once covered England but which was deforested from the Bronze Age onwards^17^.

Burying beetles (*Nicrophorus* spp.) breed on small dead vertebrates, such as rodents, and it is common for several *Nicrophorus* species to co-exist within the same woodland. Competition for carrion partitions the carrion niche so that larger burying beetle species tend to breed on larger carrion, while the smallest species (*N. vespilloides* in the UK) typically uses the smallest carrion^18–21^. Burying beetles match the number of eggs they lay to the size of the carcass they obtain^22^, and facultatively cull any surplus larvae through partial filial cannibalism^23^. The adaptive clutch size and brood size therefore depend on the size of the carrion that is routinely available for reproduction.

Here we show that the burying beetle guilds differ between the Gamlingay and Waresley Woods, and that this changes the range in carrion size available for *N. vespilloides* to breed upon. We demonstrate corresponding adaptive divergence in levels of fecundity shown by *N. vespilloides* from each population, and show how plasticity has facilitated such rapid adaptive divergence on such an ultra-local scale.

## Results

### The Nicrophorus guild differs between Gamlingay and Waresley Woods

In general, we found that the two woodlands harboured a similar number of *Nicrophorus* beetles: we caught a total of 1873 *Nicrophorus* individuals in Gamlingay Wood over the four-year sampling period compared with 1806 *Nicrophorus* individuals in Waresley Wood. However, whereas Gamlingay Wood routinely supports four *Nicrophorus* species, Waresley Wood is routinely inhabited by only two species (Fig. 1a). The average abundance per trap of each *Nicrophorus* species differed between the two woods (species × woodland interaction: *χ*^*2*^ = 142.68, d.f. = 4, *p* < 0.001). *N. vespilloides* was by far the most abundant in each woodland, comprising 80.6% (1510 individuals) of all *Nicrophorus* beetles trapped in Gamlingay Wood and 93.9% (1695 individuals) of those trapped in Waresley Wood. Both sites also contained stable populations of the largest species, *N. humator* (Fig. 1b; Supplementary Tables 1 and Supplementary Tables 2). Only Gamlingay Wood contained populations of intermediate-sized *N. interruptus* and *N. investigator* in all four years of the study (Fig. 1b; Supplementary Tables 1 and Supplementary Tables 2), and in significantly greater abundance in Waresley Woods (Tukey HSD, *z* = 7.90, *p* < 0.001 and *z* = 5.80, *p* < 0.001, respectively). The *Nicrophorus* guild is therefore significantly different between the two woodlands (PERMANOVA (permutational multivariate analysis of variance) test *F* = 0.024, *p* < 0.001; Supplementary Fig. 1b).

**Figure 1.**
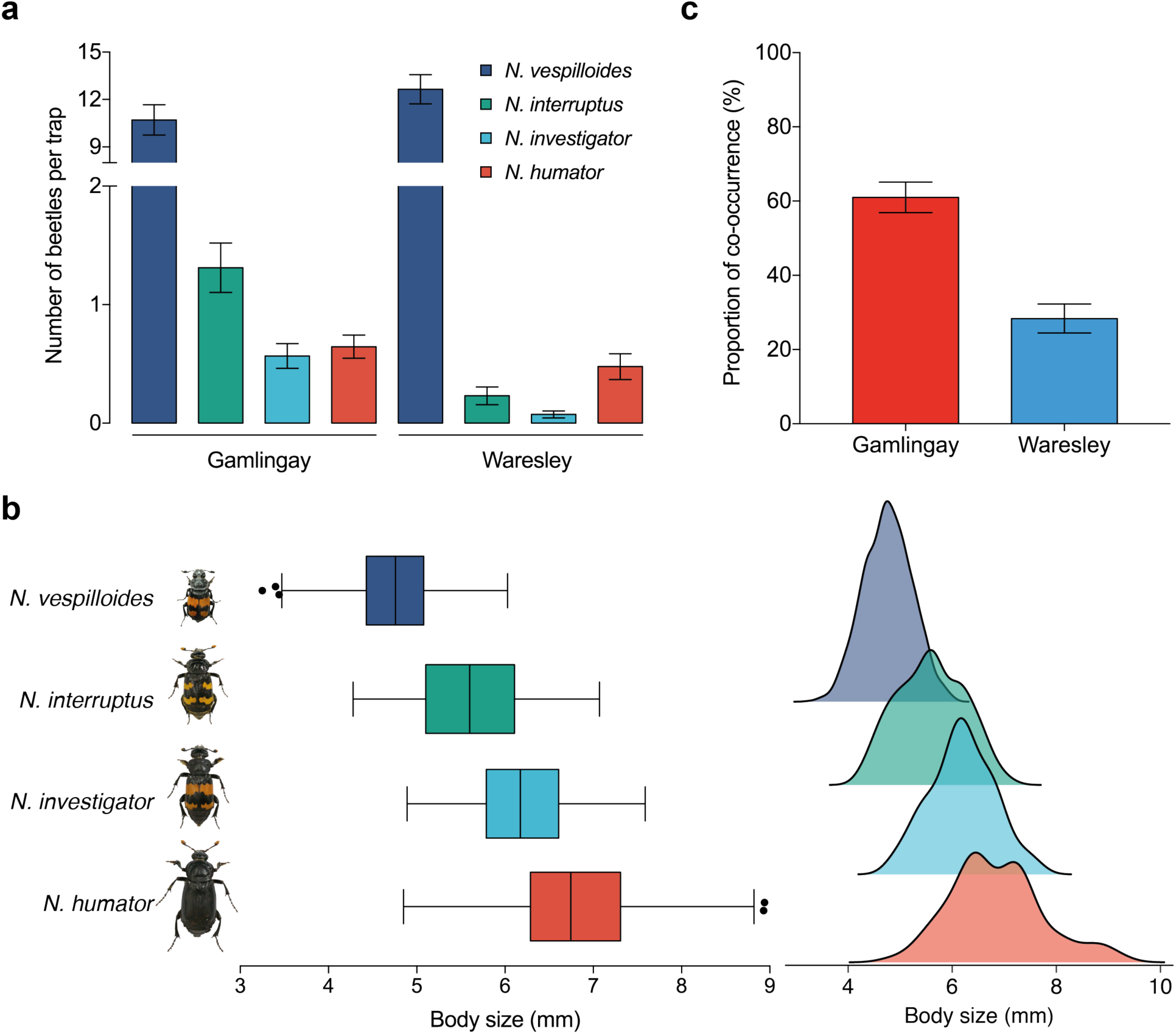
**a**, Number of *Nicrophorus* beetles caught per trap from 2014-2017. **b**, Body size and frequency distribution of field-caught *Nicrophorus* spp., illustrated withbox-and-whisker plots (left) and kernel density estimation (right). Points indicate outliers. **c**, The proportion of traps set at which *N. vespilloides* was trapped with another *Nicrophorus* species, in the two woodlands. Error bars indicate mean ± S.E.M.

### No evidence for divergence in the small mammal population between Gamlingay and Waresley Woods

We sampled the small mammal population in Gamlingay and Waresley Woods to estimate the abundance and type of carrion that might be available to the burying beetles to breed upon (see Methods). In Gamlingay, 32 animals were caught across 50 trap sessions (23 new catches and 9 recaptures); in Waresley 41 animals were caught across 50 trap sessions (30 new catches and 11 recaptures). Across both woods, bank voles (*Myodes glareolus*; range: 15-40 g) and wood mice (*Apodemus sylvaticus*; range: 13-27 g) were the dominant species, constituting 53% and 43% of all trapped mammals respectively. There was no difference in the mean body mass of small mammals sampled between the two sites (*χ*^*2*^ = 0.19, d.f. = 1, *p* = 0.661; Supplementary Fig. 2).

### The Gamlingay guild is ‘ancestral’, whereas the Waresley guild is derived

We conclude from these data that there are approximately similar numbers of *Nicrophorus* beetles within each woodland competing for an approximately similar size, abundance and type of rodent carrion to breed upon. The key difference lies in the number of burying beetle species in each wood.

We further infer from the ecological data that the more speciose burying beetle guild in Gamlingay Wood more closely approximates the ancestral burying beetle guild that was present in the ‘Wild Wood’, whereas the guild in Waresley Wood represents a more recent, derived condition. This is because burying beetle guilds in pristine ancient forests in North America are more speciose than those present in degraded woodlands^24^, in keeping with the general observation that there is a positive relationship between habitat size and species richness^25^. Furthermore, smaller-bodied generalist carrion beetles are more likely to survive in fragmented forests^24^.

### Niche expansion by N. vespilloides beetles in Waresley Wood

The ecological data additionally suggest that in Gamlingay Wood, *N. vespilloides* is more likely to be confined to breeding only on smaller carrion. In general, larger *Nicrophorus* species appear to be under selection to breed on larger carcasses^26,27^, forcing smaller beetles to breed on smaller carrion. Accordingly, *N. vespilloides* is more than twice as likely to be found on small carcasses than on large carcasses in continental forests, which are rich in *Nicrophorus* species^21^. In Waresley Wood, by contrast, *N. vespilloides* is more likely to breed on larger carrion as well because it more rarely faces competition from *N. interruptus* and *N. investigator* for larger carcasses (woodland effect: *χ*^*2*^ = 27.75, d.f. = 1, *p* < 0.001; Fig. 1c; see also ref. ^18^).

By measuring reproductive performance in the laboratory, we determined the carrion niche occupied by the following species, in decreasing order of size (Fig. 1b): *N. investigator* (from Gamlingay Wood), *N. interruptus* (from Gamlingay Wood), and *N. vespilloides* beetles (from both Gamlingay and Waresley Woods) (see Methods). For each species, we experimentally varied the size of carrion available for reproduction, presenting pairs with either a small mouse carcass (range: 12-20 g; mean ± S.E.M: 16.80 ± 0.55 g) or a large mouse carcass (range: 25-31 g; mean ± S.E.M: 28.27 ± 0.53 g; natural carcass range = 8.5-41 g). To quantify reproductive performance on each carcass size, we measured ‘carcass use efficiency’, calculated by dividing the total brood mass at the end of larval development (measured when larvae had dispersed from the carcass) by the size of the carcass the brood was reared on. We predicted that if there is local adaptation, *N. investigator* and *N. interruptus* should each exhibit greatest efficiency when breeding on a large carcass, while *N. vespilloides* from Gamlingay Wood should exhibit greatest efficiency when breeding on a small carcass. *N. vespilloides* from Waresley Wood should be more efficient at breeding on a large carcass than *N. vespilloides* from Gamlingay Wood.

We found that the efficiency of converting the carcass into larvae varied with carcass size, but in a different way for each *Nicrophorus* species (carcass size × *Nicrophorus* species interaction term: *χ*^*2*^ = 28.85, d.f. = 3, *p* < 0.001; Fig. 2 and Supplementary Table 3). *N. investigator* exhibited greater efficiency when breeding on large carcasses rather than on small carcasses (*t* = 3.51, *p* < 0.001). *N. interruptus* was similarly efficient when breeding on both large and small carcasses (*t* = 0.99, *p* = 0.325). These species are therefore adapted to breed on larger carrion. In contrast, *N. vespilloides* from Gamlingay Wood exhibited greatest reproductive efficiency on small carcasses (*t* = −4.16, *p* < 0.001). Waresley beetles were similarly efficient to Gamlingay beetles when breeding on a small carcass (*t* = 0.58, *p* = 0.938) but were significantly more efficient at breeding on a large carcass than *N. vespilloides* from Gamlingay Wood (*t* = −3.11, *p* = 0.011). We conclude that the population of *N. vespilloides* in Waresley Wood has adaptively expanded its carrion niche to breed on larger carcasses, in response to the loss of competition for larger carrion from *N. interruptus* and *N. investigator*.

**Figure 2.**
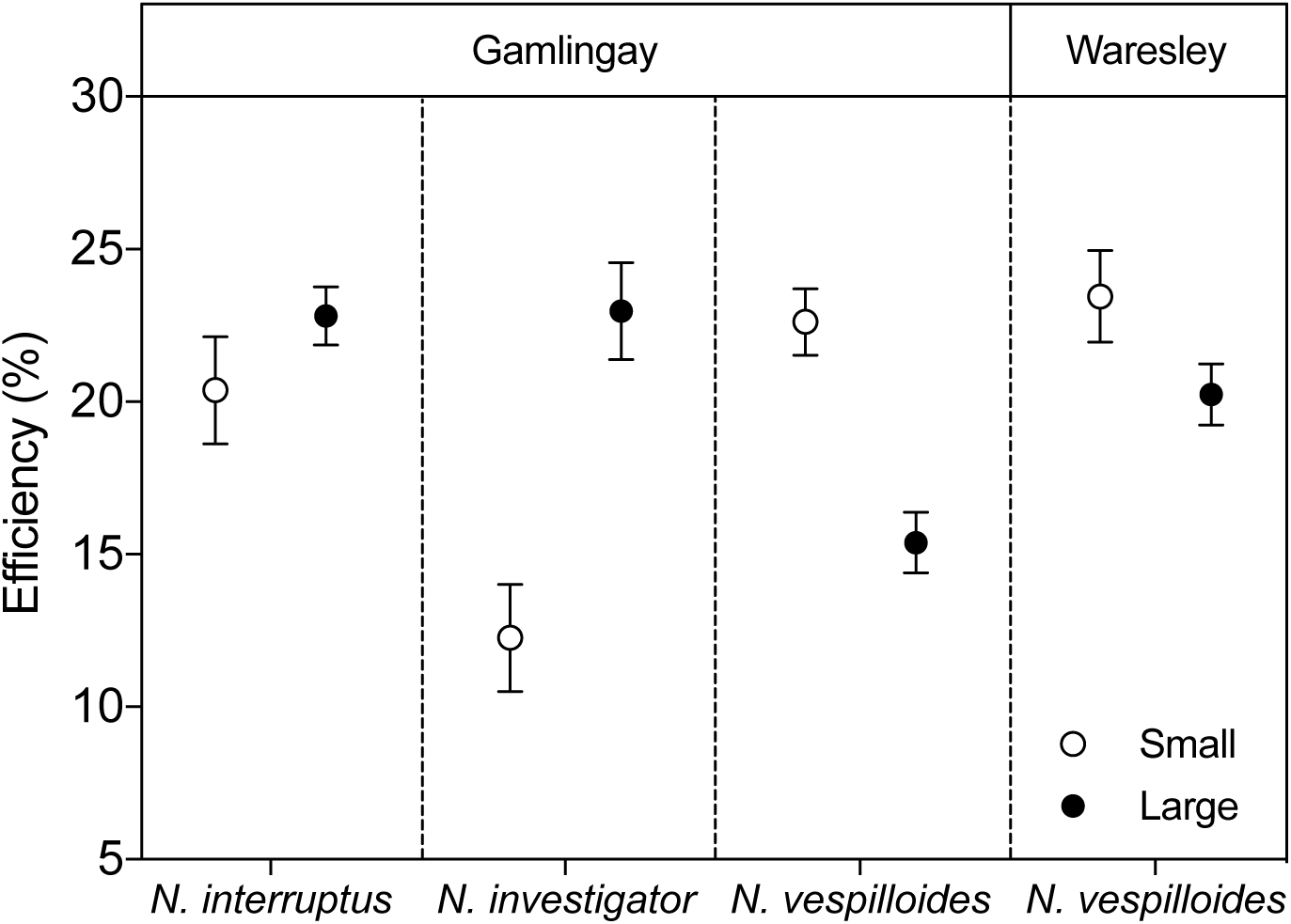
Efficiency (%) of carcass use (total brood mass divided by carcass mass) of *N. interruptus, N. investigator*, and *N. vespilloides* from Gamlingay Wood and *N. vespilloides* from Waresley Wood. Values represent the mean ± S.E.M.

### Niche expansion by Waresley N. vespilloides is due to divergent reaction norms

To test whether niche expansion by Waresley *N. vespilloides* was facilitated by genetic accommodation, we generated reaction norms relating carcass size to clutch size for *N. vespilloides* from the two woodland populations (see Methods). From the results in Fig. 2, we predicted that the reaction norm for Waresley *N. vespilloides* would be significantly steeper than the reaction norm for Gamlingay *N. vespilloides*. In fact, we found that the reaction norms differed in their intercept rather than in their slope (Fig. 3a). Although female *N. vespilloides* from both woodlands laid more eggs when given a larger carcass to breed upon (carcass size effect: *χ*^*2*^ = 10.97, d.f. = 1, *p* < 0.001; Supplementary Table 4), Waresley *N. vespilloides* consistently laid more eggs than Gamlingay *N. vespilloides* (woodland effect: *χ*^*2*^ = 21.07, d.f. = 1, *p* < 0.001; Supplementary Table 4), irrespective of carcass size (carcass size x woodland interaction: *χ*^*2*^ = 1.44, d.f. = 1, *p* = 0.231) or female size (woodland x female size interaction: *χ*^*2*^ = 1.30, d.f. = 1, *p* = 0.254).

**Figure 3.**
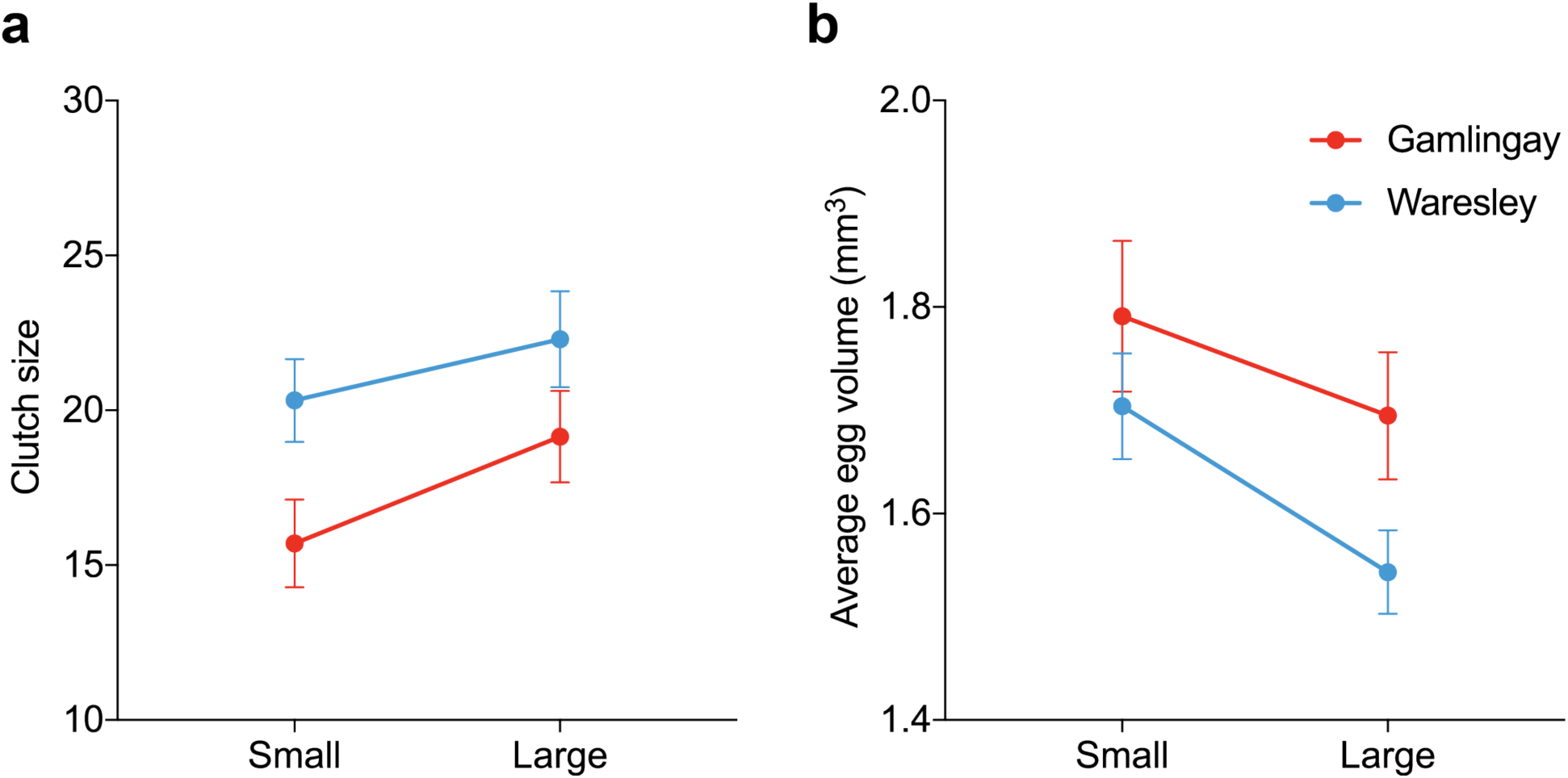
The effect of carcass size on (**a**) clutch size and (**b**) average egg volume produced by *N. vespilloides. n* = 27 Gamlingay *N. vespilloides* per carcass size treatment, and *n* = 37 Waresley *N. vespilloides* per carcass size treatment. Values represent the mean ± S.E.M.

Likewise, although female *N. vespilloides* from both woodlands laid smaller eggs when given a larger carcass to breed upon (carcass size effect: *χ*^*2*^ = 5.90, d.f. = 1, *p* = 0.015; Fig. 3b), Waresley *N. vespilloides* consistently laid smaller eggs than Gamlingay *N. vespilloides* (woodland effect: *χ*^*2*^ = 4.59, d.f. = 1, *p* = 0.032), irrespective of carcass size (carcass size x woodland interaction: *χ*^*2*^ = 0.33, d.f. = 1, *p* = 0.565). Mean egg volume per clutch did not predict mean larval mass per brood at dispersal (*χ*^*2*^ = 0.28, d.f. = 1, *p* = 0.595), regardless of the population of origin. This suggests that any under-provisioning of eggs is compensated by the over-abundance of resources available on the carcass after hatching^28,29^.

### Genetic accommodation is facilitated by plastic secondary culling of surplus offspring

When we related brood size (measured at the end of reproduction) to carcass size, we found the predicted difference between Gamlingay and Waresley *N. vespilloides* in the slope of this reaction norm (woodland x carcass size interaction: *χ*^*2*^ = 4.80, d.f. = 1, *p* = 0.029; Fig. 4; Supplementary Table 4). On a large carcass, Waresley *N. vespilloides* produced more larvae than Gamlingay *N. vespilloides* (*z* = −1.97, *p* = 0.049), whereas on a small carcass the number of larvae produced did not differ between the two populations (*z* = 0.76, *p* = 0.447). Therefore Waresley *N. vespilloides* lays more eggs on a small carcass than Gamlingay *N. vespilloides*, but there is plastic culling of surplus offspring after egg-laying, probably due to partial filial cannibalism^23^.

**Figure 4.**
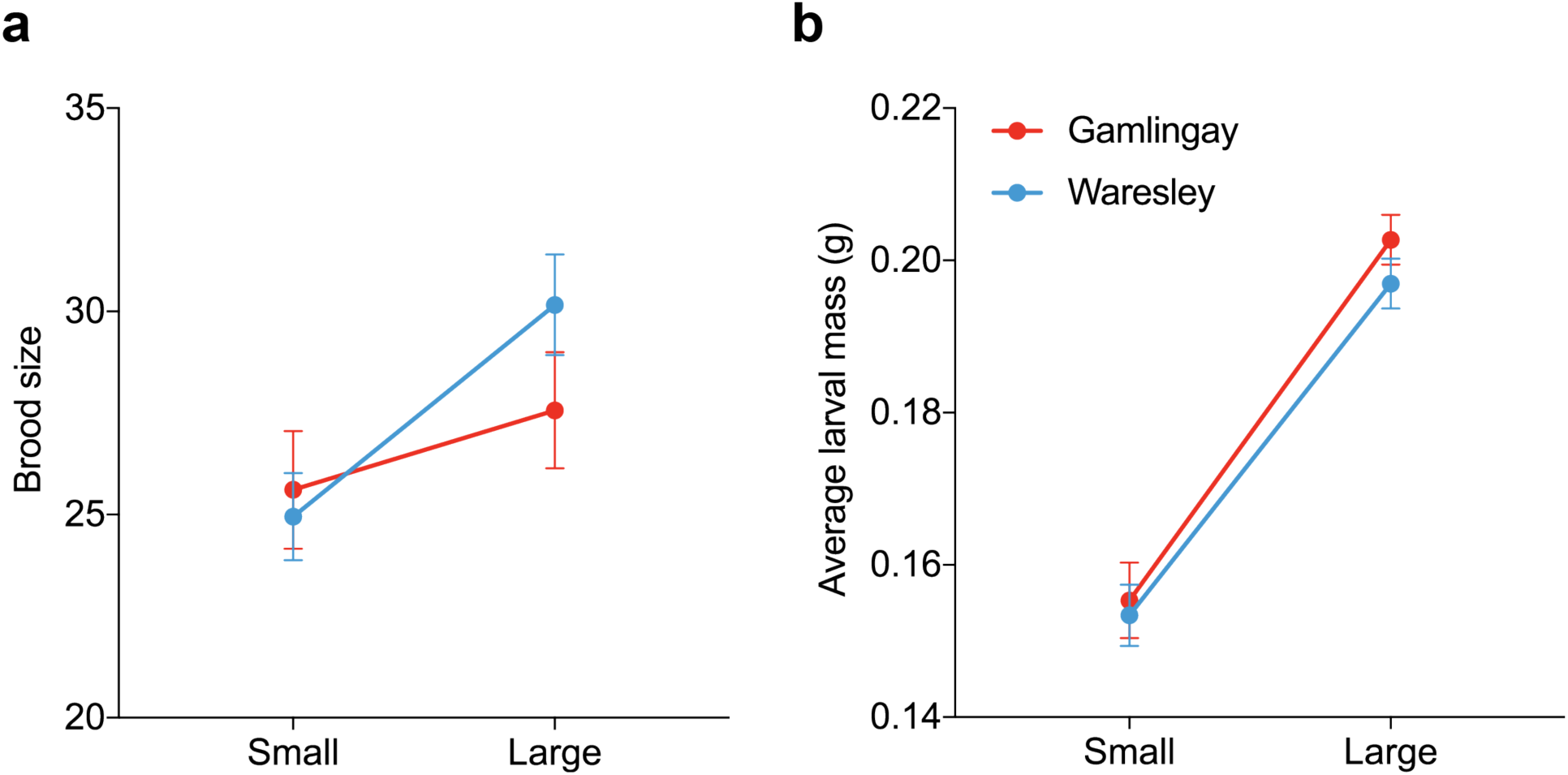
The effect of carcass size on (**a**) brood size and (**b**) average larval mass for *N. vespilloides* in the reaction norm experiment (*n* = 46 Gamlingay *N. vespilloides* per carcass size treatment, and *n* = 62 Waresley *N. vespilloides* per carcass size treatment). Values represent the mean ± S.E.M.

In short, Waresley *N. vespilloides* has evolved the ability to lay extra eggs whatever the size of the carcass it locates for reproduction. This brings fitness gains if the beetle locates a large carcass and correspondingly increases the efficiency with it converts carrion into offspring. Yet it brings fitness costs if the beetle instead finds a small carcass to breed upon, because there is a pronounced trade-off between larval number and larval size^30^ and small larvae have markedly lower fitness^31^. However, with a secondary mechanism for dispensing with excess young on smaller carcasses (probably filial cannibalism), these costs are eliminated. We suggest that genetic accommodation of the reaction norm relating clutch size to carrion has evolved and is adaptive only because there is an accompanying plastic mechanism for correcting the over-production of offspring on a small carcass. The same mechanism can also prevent costly excess fecundity if there is gene flow from Waresley to Gamlingay.

### Divergence at loci associated with oogenesis in N. vespilloides from Gamlingay v. Waresley Woods

To be certain that differences in the elevation of the reaction norm were due to evolutionary change, we sought evidence of associated genetic divergence. We generated low-coverage whole genome sequences for 40 individuals collected from each wood (*n* = 80 chromosomes per population). We found minimal divergence across the genome with no instances of extreme outliers (Supplementary Fig. 4), probably because the divergent traits are controlled by many loci. This is typical for behavioural and life history traits and consistent with the predicted quantitative genetic basis of genetic accommodation^6^. The highest *Fst*-windows in the genome showed only modest absolute values of divergence. Nevertheless, they were extreme outliers due to the otherwise consistently low pattern of *Fst* between populations (see Methods).

By comparing differentiation between both woodlands and a third population from 300km away in Swansea, Wales, UK, we polarised the divergence between populations. We calculated all pairwise *Fst* and assigned the relative divergence of each population using population branch statistics (PBS) to understand which population was driving divergence across the genome^32,33^. Not surprisingly, the distant Welsh population showed the highest genome-wide PBS. We found that population from Waresley Wood showed higher differentiation compared to the population from Gamlingay Wood (mean PBS: Waresley = 0.0074, Gamlingay = 0.0056, Wales = 0.0109). These analyses therefore support the ecological data in indicating that the Gamlingay *N. vespilloides* represents the ancestral condition, whereas the Waresley *N. vespilloides* are the derived population (Supplementary Fig. 5).

To visualise the relative divergence between populations across the genome, we generated a scatterplot of the PBS values for 2kb non-overlapping windows for each population (Fig. 5a). The analysis highlighted multiple potential candidate genes associated with the differences in egg-laying behaviour, again consistent with this trait being controlled by many loci of small effect. For example, homologs of three of the highly differentiated genes in the Waresley Wood population – *obg-like ATPase, transmembrane protein 214*, and *liprin-alpha* – show elevated expression in the ovaries of fruit flies^34^ and other arthropods^35^, suggesting a plausible role in regulating egg production. Another gene, *kekkon1*, is a transmembrane protein known to regulate the activity of the epidermal growth factor receptor (EGFR) during oogenesis in *Drosophila*^36,37^.

**Figure 5.**
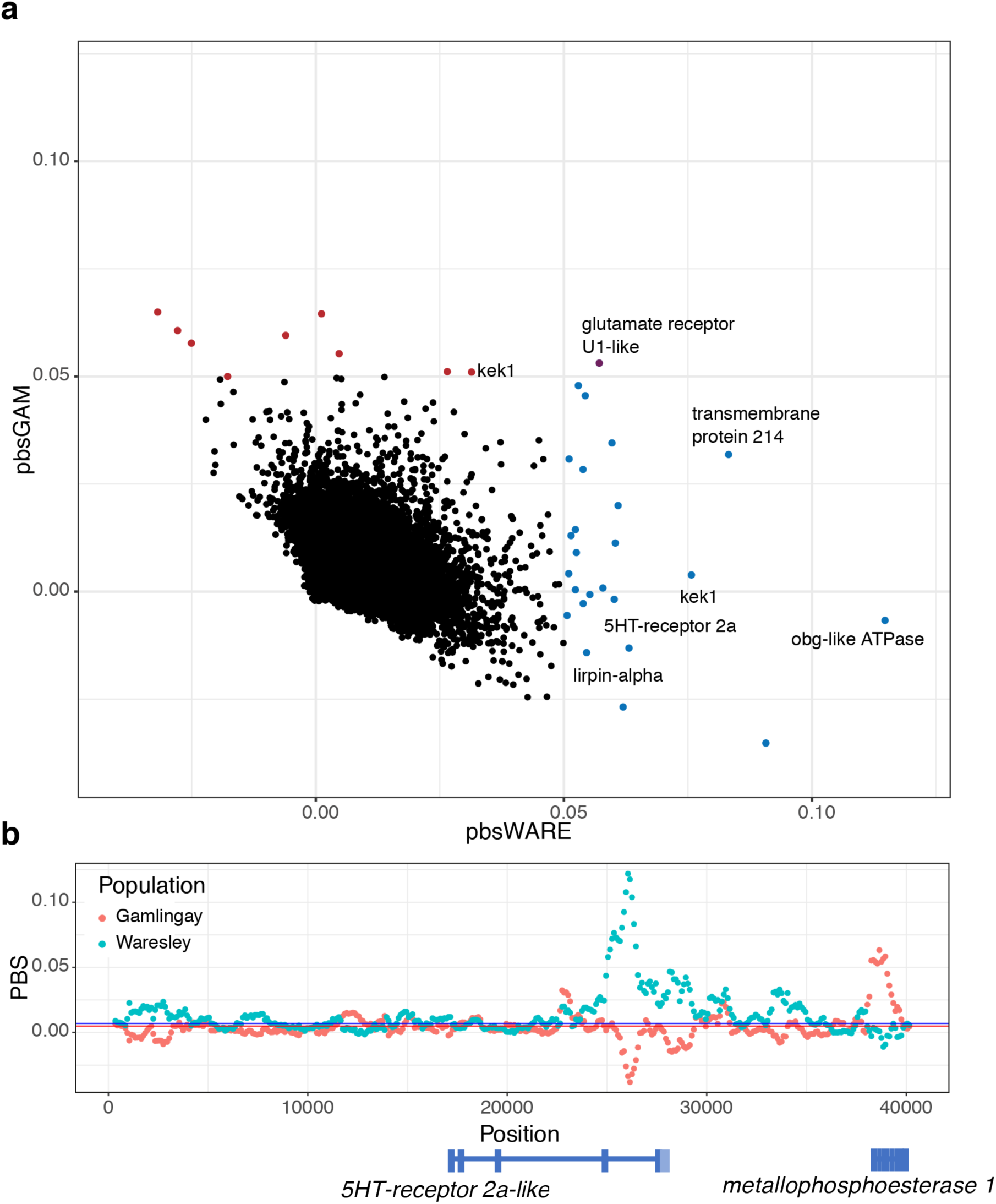
Differentiation at putative oogenesis genes. **a**, A scatterplot of PBS values for Waresley and Gamlingay in 2kb windows genome-wide. Loci in the lower right hand of the figure show high differentiation in Waresley but not in Gamlingay. Loci with PBS scores greater than 0.05 are highlighted – Waresley = blue, Gamlingay = red, Both = purple. Notable genes are annotated. **b**, Sliding window analysis (window = 500bp, slide = 100bp) of PBS values at the *5HT receptor 2a-like* receptor. The peak PBS in Waresley (blue) falls in the last intron of the gene.

Finally, we asked whether genes involved in oogenesis generally showed elevated levels of differentiation in each population, in comparison with the rest of the genome. For each population, we ranked genes by the highest PBS score in 500bp windows overlapping with the gene-body and conducted a gene set enrichment analysis for each population. *N. vespilloides* from both Waresley and Gamlingay Woods showed enrichment in multiple GO terms associated with ovaries and oogenesis (Table 1).

**Table 1.**
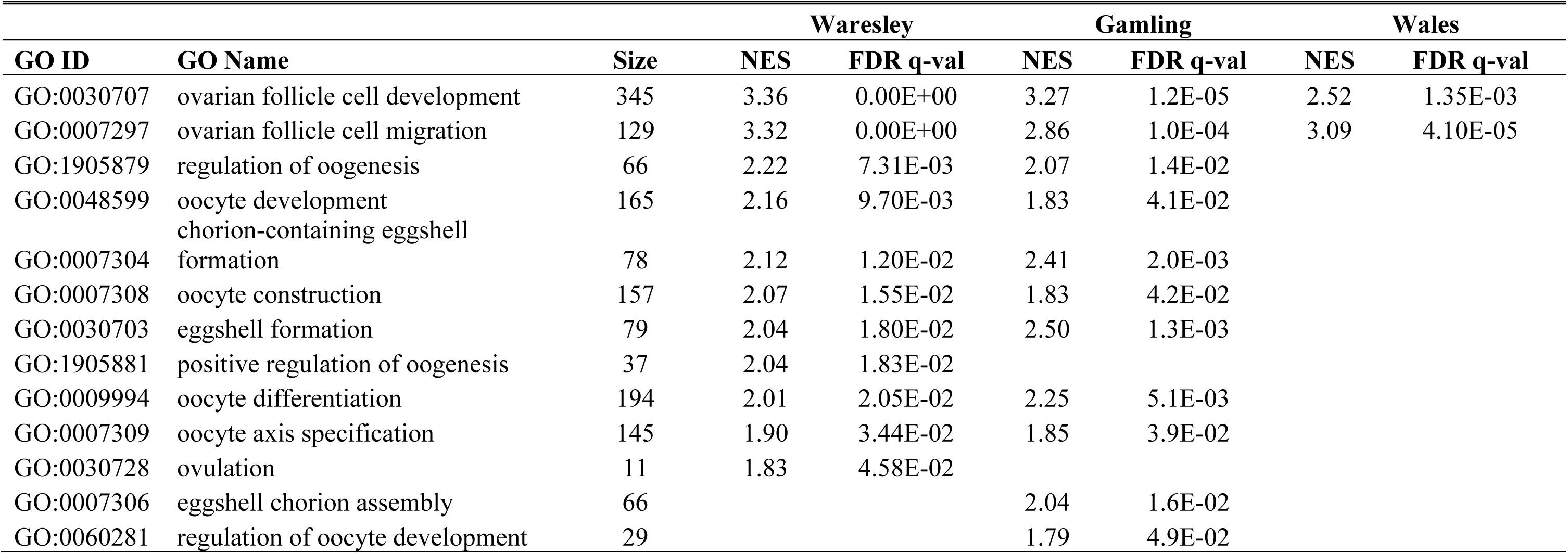
Population differences in enrichment scores in multiple GO terms associated with ovaries and oogenesis.

## Discussion

Our analyses support the three principle criteria required to show evidence for plasticity-first evolution^6^. We show that the ancestral reaction norm linking carrion size to clutch size has evolved to have a greater intercept in *N. vespilloides* from Waresley Wood. This change has enabled Waresley N. *vespilloides* beetles to expand their carrion niche in the absence of rival congenerics. It is adaptive because Waresley *N. vespilloides* are now more effective than Gamlingay *N. vespilloides* at converting carrion that ranges widely in size into offspring. Whereas previous analyses of plasticity-first evolution in wild animal populations have emphasised how new adaptations can evolve from the genetic assimilation of plastic traits^6–8,38^, our study is different in showing how adaptive traits in nature can also result from genetic accommodation.

Importantly, plasticity not only precedes the evolution of new local adaptations but also provides a secondary correcting mechanism that ameliorates the costs of expressing these new adaptations in the wrong local environment. Indeed, the presence of a plastic secondary correcting mechanism is key to explaining how there has been genetic accommodation of the reaction norm in *N. vespilloides* from Waresley Woods, despite gene flow with *N. vespilloides* from Gamlingay Woods. We have demonstrated that when evolution by genetic accommodation in one trait is coupled with a second plastically expressed correcting mechanism, then new adaptations can emerge rapidly and on an ultra-fine scale. We suggest that whenever these two mechanisms operate together, in two related traits, there is immense potential for rapid evolutionary change that is finely tuned to local environmental conditions.

## Methods

### The history of Gamlingay and Waresley Woods

We focused on two woodlands: Gamlingay Wood (Latitude: 52.15555°; Longitude: −0.19286°), and Waresley Wood (Latitude: 52.17487°; Longitude: −0.17354°). They are woodland islands of approximately the same size (c 50ha) in a landscape dominated by arable farming (Supplementary Fig. 1a). In common with other woodlands recorded in the Domesday book, these woods have stayed approximately the same size since 1086^39^. Gamlingay Wood was acquired and managed by Merton College, Oxford for c. 800 years^40^. Its ecology was described in detail in 1912^40^. The modern history of Waresley Wood is less well-known^39,41^. Both sites are now designated as ‘ancient woodland’ and are managed by the Bedfordshire, Cambridgeshire and Northamptonshire Wildlife Trusts.

### Competition for carrion in Gamlingay and Waresley Woods

#### Burying beetle trapping

Each year from 2014-2017 we set five beetle traps per woodland, at exactly the same five locations within each wood (Supplementary Fig. 1a), and checked the contents every 2-3 weeks from June until October, each time rebaiting the trap with fresh compost and a dead mouse. Carrion-baited soil-filled traps were suspended at each site, with traps set at least 150 m apart from each other. We collected and identifed all the *Nicrophorus* spp. within each trap and measured the pronotum width (to the nearest 0.01 mm) as an index of body size (see below).

We caught five species in total (in increasing order of size): *N. vespilloides, N. interruptus, N. vespillo, N. investigator* and *N. humator*. However, *N. vespillo* were only caught 5 and 3 times within the four years in Gamlingay and Waresley respectively, indicating that there is no stable population of *N. vespillo* in either wood. A clear separation of the guild structure between the woods can be seen for each sampling time across the four years in NMDS (nonmetric multidimensional scaling) ordination (Supplementary Fig. 1b). Evidence from other populations suggests that our measurements reflect long-term differences in guild structure because abundance measures of *Nicrophorus* are robust over time, even when there are marked change in woodland management^40^.

#### Burying beetle husbandry in the lab

After removing any phoretic mites, beetles were retained and kept individually in plastic boxes (12cm × 8cm × 2cm), which were filled with moist soil in a laboratory kept at 20°C and on a 16:8 light to dark cycle. Beetles were fed twice a week with minced beef. We kept all field-caught individuals for at least two weeks before breeding to ensure that they were sexually mature and to reduce any variation in nutritional status. We then maintained stock populations of both Gamlingay and Waresley Woods by breeding pairs of unrelated individuals on 8-16 g mice carcasses.

#### Size distributions of the Nicrophorus spp

Body size was measured for *Nicrophorus* spp. collected from 2014 to 2017 (except in 2016). In total, 838 *N. vespilloides*, 41 *N. humator*, 127 *N. interruptus*, 54 *N. investigator*, and 5 *N. vespillo* were measured for Gamlingay Wood, whereas 824 *N. vespilloides*, 51 *N. humator*, 25 *N. interruptus*, 4 *N.* investigator, and 6 *N. vespillo* were measured for Waresley Wood. Mean body size of *Nicrophorus* spp. significantly varied among species (GLMM: *χ*^*2*^ = 2069.44, d.f. = 4, *p* < 0.001; Supplementary Table 2). Post-hoc comparisons revealed that *N. vespilloides* was smaller than the other *Nicrophorus* spp. (Supplementary Table 1 and Supplementary Table 2). A Kolmogorov-Smirnov test comparing pairwise differences in *Nicrophorus* spp. body size frequency distribution revealed similar patterns found in differences of mean body size (Fig. 1 and Supplementary Table 5).

#### Mark-recapture experiment

In 2014, we investigated the interconnectivity of populations of burying beetles between Gamlingay and Waresley Woods using a mark-recapture survey. In total, 98 *N. vespilloides*, 9 *N. humator*, 17 *N. interruptus*, 9 *N. investigator,* and 2 *N. vespillo* were marked for Gamlingay Wood, whereas 113 *N. vespilloides*, 5 *N. humator*, 1 *N. interruptus*, 1 *N. investigator* were marked for Waresley Wood. Beetles were marked with a numbered plastic bee tag on either the right or left elytra for those found in Gamlingay or Waresley Woods respectively. To identify any previously caught beetles that lost their tags, we permanently marked them by cutting a small portion of the elytra (less than 2%). All marked beetles from each wood were released from a designated location at the geographic midpoint with the minimum total distance to all trapping sites for Gamlingay (Latitude: 52.16294°; Longitude: −0.18984°) and Waresley Woods (Latitude: 52.176508°; Longitude: −0.156776°). We found no evidence of migration between the two sites. We recaught 8 of 98 marked *N. vespilloides* from Gamlingay Wood in Gamlingay Wood and 8 out of 113 marked *N. vespilloides* from Waresley Wood in Waresley Wood. None of the other marked *Nicrophorus* spp. was recaptured in either wood.

#### Small mammal trapping

To assess the rodent carrion available for *Nicrophorus* spp. reproduction, we sampled the small mammal communities in the two woodlands. In general, rodent populations peak in the autumn, because breeding for the year has just ceased and there has yet to be any winter-induced mortality^42^. Sampling at this time is therefore ideal for detecting which species are present and for determining their relative abundance. We placed Longworth traps in both woodlands in November 2016. Traps were baited with oats and blowfly maggots (with hay provided as bedding) and set in pairs within 20 m of each original beetle trapping site (Supplementary Fig. 1a), with 10 traps set per wood. We continuously trapped rodents for three days, generating 50 trap sessions per woodland. Traps were checked daily at approximately 0830 and 1500 (generating a total of 30 trap sessions overnight and 20 trap sessions in daylight hours). Trapped mammals were identified, weighed, sexed, marked by a fur clip on either the right or left rear flank, and released *in situ*. Any recaptured mammal was recorded in subsequent censuses. All traps were reset and rebaited immediately after checking. In addition to the results reported in the main text, one yellow-necked mouse (*Apodemus flavicollis*; range: 14-45 g) and one common shrew (*Sorex araneus*; range: 5-14 g) were caught in Waresley. We have no reason to think that the mortality of these rodent species should differ between woodlands that are in such close proximity and that are subjected to similar levels of ecological management.

#### Division of the carrion niche by Nicrophorus beetles in Gamlingay and Waresley Woods

During the field seasons in 2017 and 2018, pairs of wild-caught *N. vespilloides, N. interruptus*, and *N. investigator* were bred on either small (12-20 g; 16.80 ± 0.55 g) or large carcasses (25-31 g; 28.27 ± 0.53 g) within a breeding box (17 × 12 × 6 cm) filled with 2 cm of moist soil. All field-caught beetles were kept for two weeks in the laboratory and fed twice a week prior to breeding. In total, we established eight treatments: large (*n* = 47) and small (*n* = 48) carcasses for Gamlingay *N. vespilloides*; large (*n* = 42) and small (*n* = 33) carcasses for Waresley *N. vespilloides*; large (*n* = 25) and small (*n* = 18) carcasses for *N. interruptus*; large (*n* = 9) and small (*n* = 13) carcasses for *N. investigator*. *N. interruptus* and *N. investigator* were both drawn from Gamlingay Wood as the populations of these species in Waresley were too small to be used experimentally.

Approximately eight days after parents are given a carcass to breed on, larvae switch from aggregating on the carcass to dispersing away into the soil to pupate. When one or more larvae from each brood switched their behaviour in this way, we scored the whole brood as having reached the dispersal stage. At this point, all larvae were counted and total brood mass was weighed to the nearest 0.001 g. We also calculated average larval mass for each brood by dividing total brood mass by number of larvae.

In addition to the results reported in the main text, we found that Gamlingay *N. vespilloides* was less efficient on larger carcasses than both *N. interruptus* (*t* = −3.92, *p* < 0.001) and *N. investigator* (*t* = −2.69, *p* = 0.038).

#### Niche expansion in N. vespilloides by genetic accommodation

This experiment was conducted in the laboratory, over two blocks in 2017, using the second and third descendant generations of field-caught beetles from Gamlingay and Waresley Woods. By rearing beetles from both woodlands in the lab in a common garden environment for at least one generation prior to testing, we minimized any residual environmental effects when quantifying the reaction norm for each population. To pair beetles for reproduction, we began by haphazardly casting broods into dyads when larvae had matured into adults. Within each dyad, we haphazardly chose four males from one brood, and paired them with four haphazardly chosen females from the second brood. Two of these pairs were then given a small mouse to breed upon (12-17 g; 15.03 ± 0.67 g), while the remaining two pairs were given a large mouse to breed upon (26-31 g; 28.86 ± 0.67 g). By using sibships to generate pairs in this way, we were able to compare how very similar genotypes responded to the opportunity to breed on either a small or large mouse. Each pair, and their mouse, was housed in a clear plastic box (17 × 12 × 6 cm), with 2 cm depth of Miracle-Gro compost. The box was placed in a dark cupboard for eight days after pairing the beetles, until larvae started to disperse away from the carcass. In the second block of the experiment, we measured clutch size. Fifty-six hours after we introduced the beetles to the carcass, we photographed the base of each transparent breeding box. Using Digimizer ver. 5.1.0, we then counted the number of visible eggs, and also measured the length (L) and width (w) of all eggs that were able to be measured accurately (i.e. those that were fully visible and lying flat on the base of the box). All counting and measuring of eggs was performed blind to the carcass size treatment and the population from whence the breeding beetles came. Egg volume was then calculated using the formula V=1/6 *π*w^2^*L, which assumes eggs to be a prolate spheroid (following ref. ^43^). In both blocks, we also measured the number of larvae present at dispersal and weighed the whole brood.

#### Clutch size

In total, 2518 eggs were counted across 132 breeding boxes, of which 1633 could be measured. No eggs could be seen in four of these boxes, and these were excluded from further analysis (hence *n* = 128). The total number of eggs counted (observed clutch size), and the number of eggs which could accurately be measured, each correlated positively with brood size (number of eggs counted: *χ*^*2*^ = 27.18, d.f. = 1, *p* < 0.001; number of eggs measured: *χ*^*2*^ = 24.26, d.f. = 1, *p* < 0.001), indicating these are accurate estimates of true clutch size. The total volume of all eggs in a clutch did not differ between populations (*χ*^*2*^ = 2.16, d.f. = 1, *p* = 0.141; Supplementary Table 4) or carcass size treatments (*χ*^*2*^ = 2.38, d.f. = 1, *p* = 0.123; Supplementary Table 4).

#### Brood size

We found that when given a larger carcass for reproduction, *N. vespilloides* from Gamlingay Wood produced only a marginally larger brood than when breeding on a small carcass (*z* = 1.82, *p* = 0.068). By contrast, Waresley *N. vespilloides* produced significantly larger broods on larger carcasses than on smaller carcasses (*z* = 5.53, *p* < 0.001). Whether they bred on small or large carcasses, the variance was similar for Gamlingay and Waresley *N. vespilloides* in both brood size (Bartlett’s test, *p* = 0.615) and carcass use efficiency (Bartlett’s test, *p* = 0.943).

#### Offspring size

Females produced heavier larvae at dispersal when breeding on a large carcass than on a small carcass (*χ*^*2*^ = 139.05, d.f. = 1, *p* < 0.001; Supplementary Table 4), irrespective of their woodland of origin (*χ*^*2*^ = 1.06, d.f. = 1, *p* = 0.304; Supplementary Table 4). We could find no difference in the size of adult *N. vespilloides* trapped in Gamlingay and Waresley Woods, (*χ*^*2*^ = 2.02, d.f. = 1, *p* = 0.156; Supplementary Fig. 3a), even after controlling for sex (*χ*^*2*^ = 1.41, d.f. = 1, *p* = 0.235). We could not detect any differences in body size distribution between Gamlingay and Waresley *N. vespilloides* (*D* = 0.044, *p* = 0.410; Supplementary Fig. 3b). Thus, given an abundance of resources for reproduction, *N. vespilloides* allocates them to producing more offspring rather than larger offspring.

#### Divergence at loci associated with oogenesis in N. vespilloides from Gamlingay v. Waresley Woods

We generated low-coverage whole genome sequences for three populations of *N. vespilloides*: from Waresley Wood, Gamlingay Wood and Swansea, Wales, UK. Twenty-two *N. vespilloides* from three ancient woodlands near Swansea in Wales were trapped by Dr Chris Cunningham in 2017. The three sites in Wales (Park Wood (Latitude: 51.57258°; Longitude: −4.03096°); Clyne Valley Wood (Latitude: 51.61262°; Longitude: −4.02293°); and Caswell Bay Wood (Latitude: 51.57258°; Longitude: −4.03096°)) are approximately 300 km away from our two study sites in Cambridgeshire.

DNA was individually extracted from beetle heads using the DNeasy Blood and Tissue kit (Qiagen) and subsequently quantified and quality checked using NanoDrop and Qubit. DNA was then shipped to Cornell University, where libraries were prepared using partial reactions of a Nexterra kit by the Cornell Genomics Core. Libraries were subsequently sequenced by Novogene (Davis, CA, USA) at an average depth of approximately 5× coverage. Paired-end 550 bp insert libraries were prepared for each sample with the Nextera library preparation kit. Libraries were sequenced using the Illumina HiSeq (Novogene, Davis, CA) at an average coverage of 3.4X. Trimmomatic (v0.36) was used to removed adaptors and poor-quality sequence. Trimmed reads were mapped to the *N. vespilloides* reference genome using the Burrows-Wheeler Aligner (v0.7.13)^44^. SNPs were identified using Picard (v2.8.2) and GATK (v3.6) HaplotypeCaller following best practice recommendations^45^. After alignment, SNPs were hard filtered using the parameters: QD < 2.0 ‖ SOR > 3.0 ‖ FS > 200. We used VCFtools to calculate population genetic statistics for each population. To examine population structure, we generated a thinned VCF file with one SNP per 5kb and used Tassel (version 5)^46^ to calculate generate an MDS plot. For calculation of *Fst* and PBS values, aligned bam files were analysed in ANGSD (v 0.911)^47^, which is specifically designed for analysis of low-coverage genome sequencing data. Data for this project is available at the NCBI Sequence Read Archive under Bioproject PRJNA530213.

Genes were assigned gene ontology (GO) terms using the BLAST2GO workflow (v5.1.1). In brief, gene identity was determined based upon BLAST searches to the Arthropod or *Drosophila* non-redundant protein databases and protein domains were identified based on matches to Interpro database. GO terms were assigned to each gene model based upon mapping results. GO terms were filtered with the “Filter Annotation by GO Taxa” option to remove GO terms that are incompatible for Arthropods.

Population genetic summary statistics were similar for both populations (mean of 2 kb windows – Gamlingay Pi = 0.0055 ± 1.02e-5, Tajima’s D = −0.70 ± 0.002; Waresley Pi = 0.0056 ± 1.0e-5, Tajima’s D = −0.67 ± 0.002). Consistent with previous microsatellite analyses^16^, we found little to no genetic differentiation between populations from Gamlingay and Waresley Woods (unweighted *Fst* = 0.0069; weighted *Fst* = 0.013), strongly suggesting there is ongoing gene flow between the two populations.

Among the few loci that diverged between the populations, the top window of divergence between the two populations fell in *transmembrane protein 214* (*Fst* = 0.11, *zFst* = 19.2, *p* = 7.2e-82), which is highly expressed in the ovaries in *Drosophila melanogaster*, and is therefore a potential candidate gene influencing differences in *N. vespilloides* egg laying behaviour between Gamlingay and Waresley Woods.

PBS analysis identified further candidate genes. We found that a different portion of *kek1* is moderately differentiated in *N. vespilloides* from Gamlingay Wood (Fig. 5a), suggesting that differentiation related to oogenesis is not limited to *N. vespilloides* from Waresley Wood. One of the stronger signals of differentiation in Waresley Wood that may contribute to the phenotypic differences observed between populations is found in a serotonin receptor (Fig. 5b). Serotonin has been linked to reproduction via effects on the production of ecdysteroids such as juvenile hormone in multiple insects^48–50^. Serotonin has also been related to the intensity of aggressive behaviour in contests across diverse insect groups^51–54^. Consistent with the outlier analyses, oogenesis-related GO-terms tended to have higher enrichment scores in *N. vespilloides* from Waresley Wood compared to Gamlingay Wood (Table 1). The gene enrichment analysis also revealed local divergence between beetles from Waresley and Gamlingay Woods at genes associated with other traits, including learning and memory and sensory systems (Supplementary Data 1).

### Statistical analyses

All statistical analyses were performed in R version 3.4.3 (R Development Core Team), with NMDS in the package *vegan*, GLM and GLMM in the package *lme4*^55^, and Tukey’s HSD post hoc comparisons in the package *lsmeans*^56^.

#### Competition for carrion in Gamlingay and Waresley Woods: field data

To test for a difference in the *Nicrophorus* guild between Gamlingay and Waresley Woods, a frequency table of beetle communities was analysed using permutational multivariate analysis of variance (PERMANOVA) on two-dimensional nonmetric multidimensional scaling (NMDS) based on Bray-Curtis distances^57^, with the ‘adonis’ function (vegan package). We tested the effect of study site on the composition of the beetle community, using sampling year as strata. The analysis was based on 10000 permutations of the data. We visualized the difference of beetle community between Gamlingay and Waresley population in two dimensions of a NMDS plot. NMDS two-dimensional stress values (a measure of goodness of fit) were below 0.1 (0.085), indicating the ordination provides a good fit to the data^58^.

We used general linear mixed models (GLMM) with a Poisson error structure to test for differences across beetle species and sites on average abundance for each species per trap. Beetle species and population (Gamlingay/Waresley) were included as fixed effects, whereas trap ID and sampling year were included as random factors. We also tested for differences in the probability of co-occurrence (co-occurring = 1, non-co-occurring = 0) between *N. vespilloides* and at least one other *Nicrophorus* spp. across sites using a binomial GLMM. Population was included as a fixed effect and year as a random effect.

We compare the distributions of body size between Gamlingay and Waresley *N. vespilloides*, and among *Nicrophorus* spp., using the Kolmogorov-Smirnov (K-S) two-sample test, which tests whether the cumulative distributions of two data sets are derived from the same distribution. We also tested for significant differences in mean body size between *N. vespilloides* in Gamlingay and Waresley, using a GLMM that included population (Gamlingay/Waresley) and sex (male/female) as fixed effects, and sampling year as a random factor. Differences of mean body size among *Nicrophorus* spp. was assessed in a GLMM that included species and sex as fixed effects, and sampling year as a random factor.

We analysed the body mass of rodents in Gamlingay and Waresley using a GLM by including rodent species and population (Gamlingay/Waresley) as fixed effects.

#### Division of the carrion niche by Nicrophorus beetles in Gamlingay and Waresley Woods

To test for differences in reproductive performance between species, we conducted a GLMM regression to analyse differences in efficiency (total brood mass divided by carcass mass), which was logit transformed prior to analysis. Beetle species, carcass size (small/large), and their interaction were included as explanatory variables. Sampling year was included as a random factor. In this analysis, we included beetle species as *N. interruptus, N. investigator*, Gamlingay *N. vespilloides* and Waresley *N. vespilloides* to fully compare differences not only between *Nicrophorus* beetle species, but also *N. vespilloides* between populations.

#### Niche expansion in N. vespilloides by genetic accommodation

For the reaction norm experiment, we used GLMMs to test the interacting effect of population and carcass size on brood size and average larval mass, with dyad identity nested within block included as a random factor. In all models, brood size and average larval mass were analysed with a Poisson and Gaussian error distribution, respectively. A similar statistical approach was used for analyses of clutch traits to test for the significant differences on clutch size and average egg volume in GLMMs with a Poisson and Gaussian error distribution, respectively. The effect of population of origin, carcass size, and their interaction were included as fixed effects, whereas dyad identity was included as a random factor. If a significant interaction was found, a Tukey’s post-hoc test was performed to detect significant effects using multiple pairwise comparisons.

## Acknowledgements

We thank P. Pilbeam for carrying out small mammal trapping in the field. We thank A. Attisano, C. Swannack, and R. Mashoodh for helping with beetle trapping. S.-J.S. was supported by the Taiwan Cambridge Scholarship from the Cambridge Commonwealth, European & International Trust. A.M.C. was supported by a studentship from the National Environmental Research Council’s Earth System Sciences Doctoral Training Partnership at Cambridge University and a Victoria Brahm Schild Internship Grant from Homerton College. R.M.K. was supported by a European Research Council Consolidator’s grant 301785 BALDWINIAN_BEETLES and a Wolfson Merit Award from the Royal Society.

## Author Contributions

R.M.K. conceived the idea. S.-J.S., A.M.C. S.P. and R.M.K designed the study. S.-J.S., A.M.C. and B.J.M.J. acquired the phenotypic data, which was analysed by S.-J.S. and A.M.C. M.J.S. and S.P. designed the genetic’s component, which was collected by S.P. and analysed by M.J.S and S.E.M. All authors contributed to discussion of the results and writing the manuscript.

## Competing Interests Statement

The authors declare no competing interests.

## Supplementary Information

**Supplementary Fig. 1.**
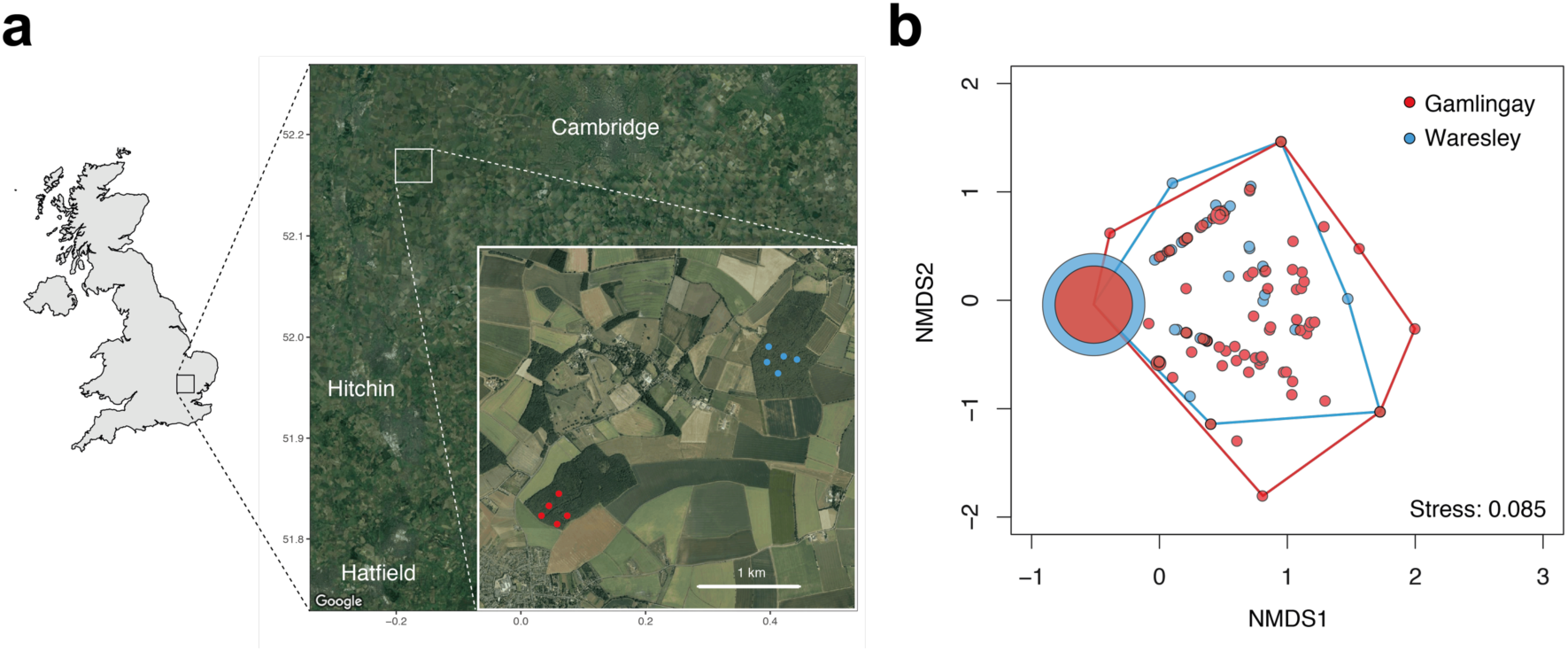
Study sites and population differences in community structure. **a**, Location of traps in Gamlingay Wood (red points) and Waresley Wood (blue points). **b**, NMDS ordination of Gamlingay (*n* = 141) and Waresley (*n* = 134) community structure. Each point represents the sum of the beetle community collected in each trap per sampling time.

**Supplementary Fig. 2.**
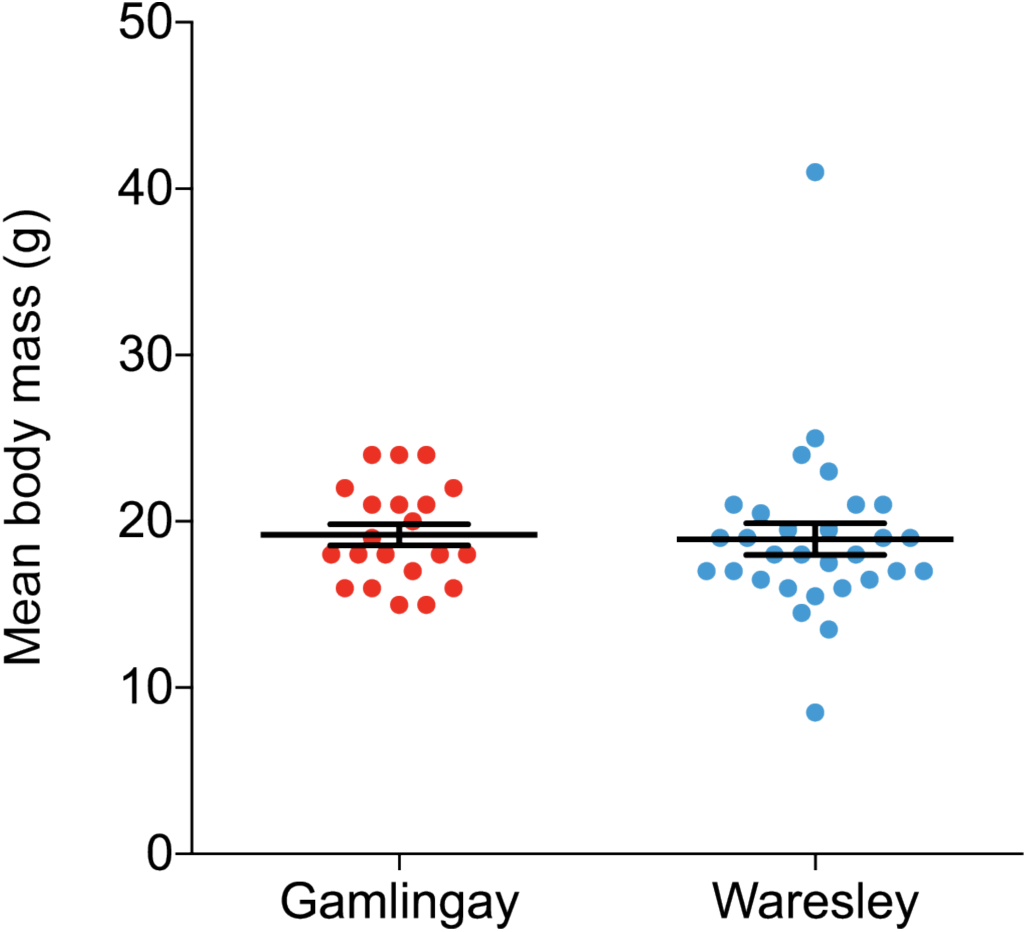
Mean body weight of small mammals. Gamlingay (*n* = 21, red) and Waresley Woods (*n* = 30, blue). One bank vole and one wood mouse, each caught in in Gamlingay Wood, escaped before they could be measured.

**Supplementary Fig. 3.**
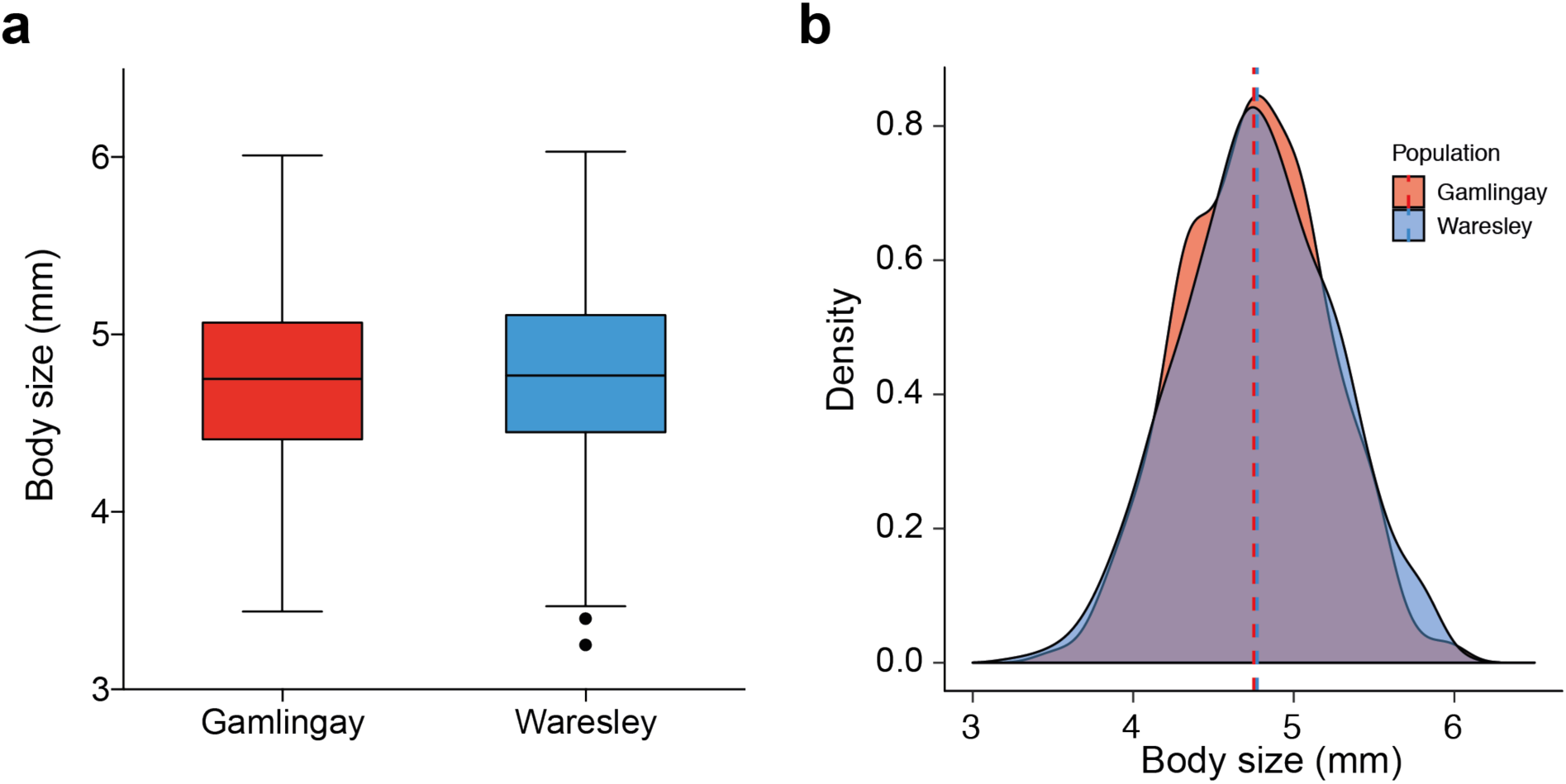
Differences in (a) body size and (b) frequency distribution of field-caught *N. vespilloides* from Gamlingay and Waresley Woods. Measurements depict pronotum width, the standard index for measuring beetle body size. Median values, inter-quartile range, maximum, and minimum are as illustrated in box-and-whisker plots. Outliers are depicted as points. Number of *N. vespilloides* sampled: Gamlingay (*n* = 839) and Waresley Woods (*n* = 824).

**Supplementary Fig. 4.**
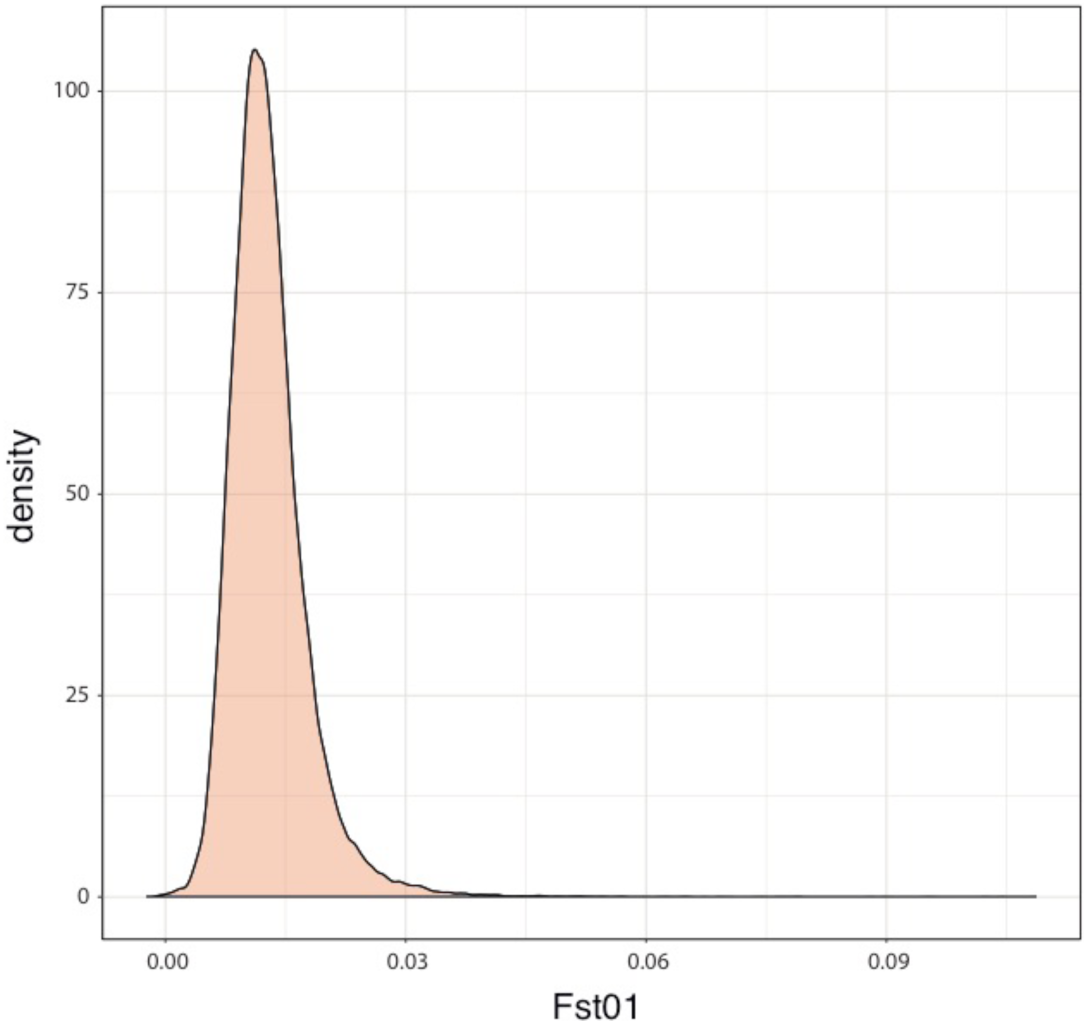
Density plot of *Fst* value between Gamlingay and Waresley Woods. The plot shows the narrow distribution of low *Fst* values between the two woodland populations. Though there is a long tail to the right, the extreme values are modest in absolute terms.

**Supplementary Fig. 5.**
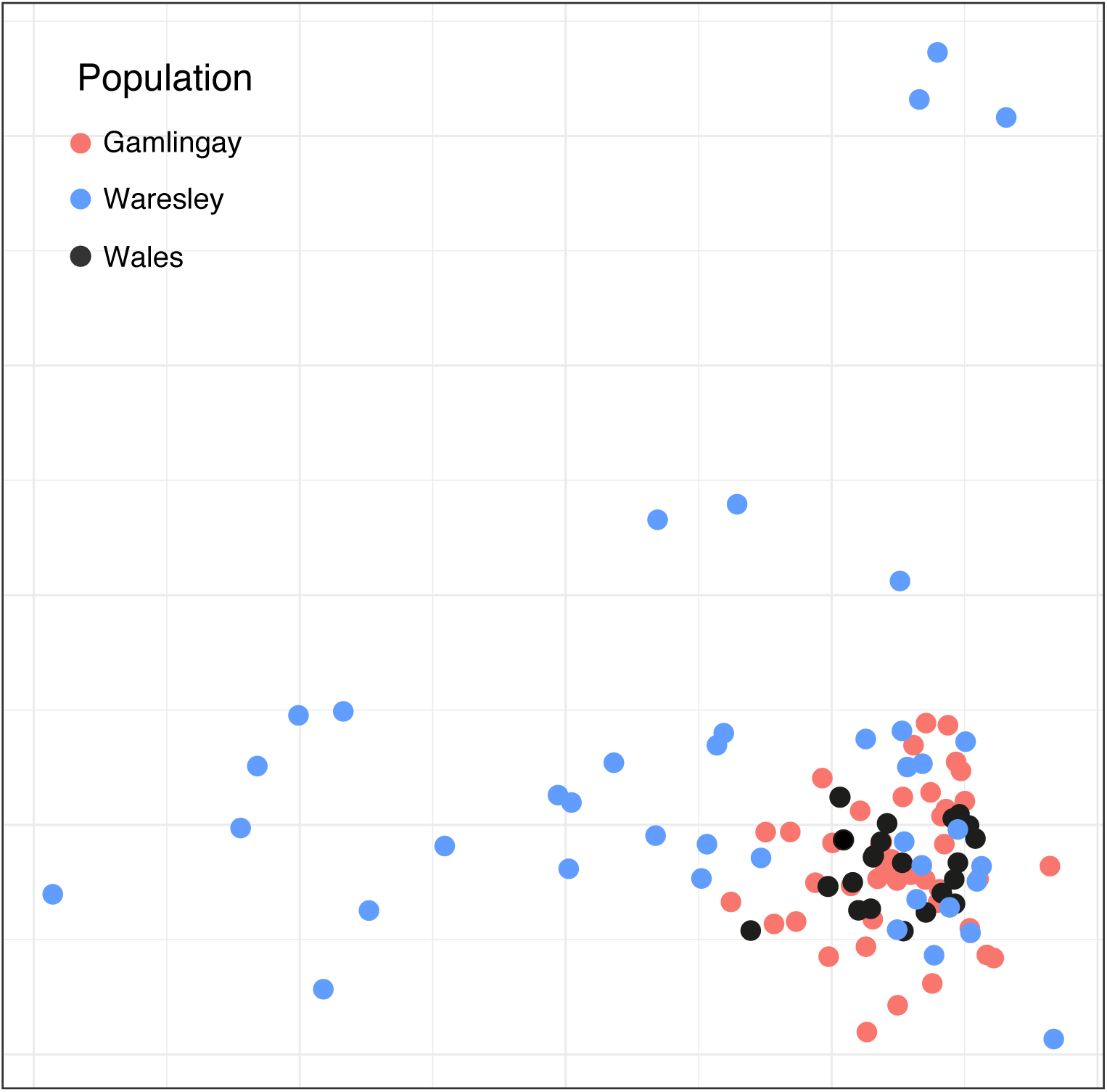
MDS plot of three burying beetle populations. The plot shows the first two dimensions of a multidimensional scaling analysis of genetic diversity among burying beetles in three populations. The populations do not separate out, indicating little to no genetic structure.

**Supplementary Table 1.**
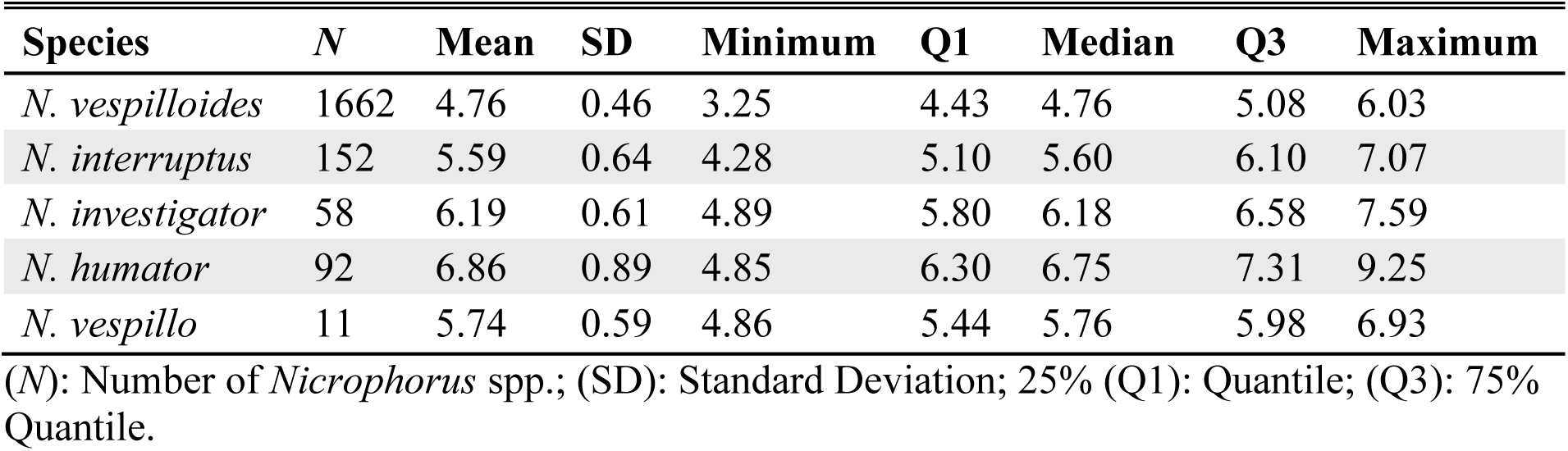
*Nicrophorus* spp. body size (given by pronotum size, in mm).

**Supplementary Table 2.**
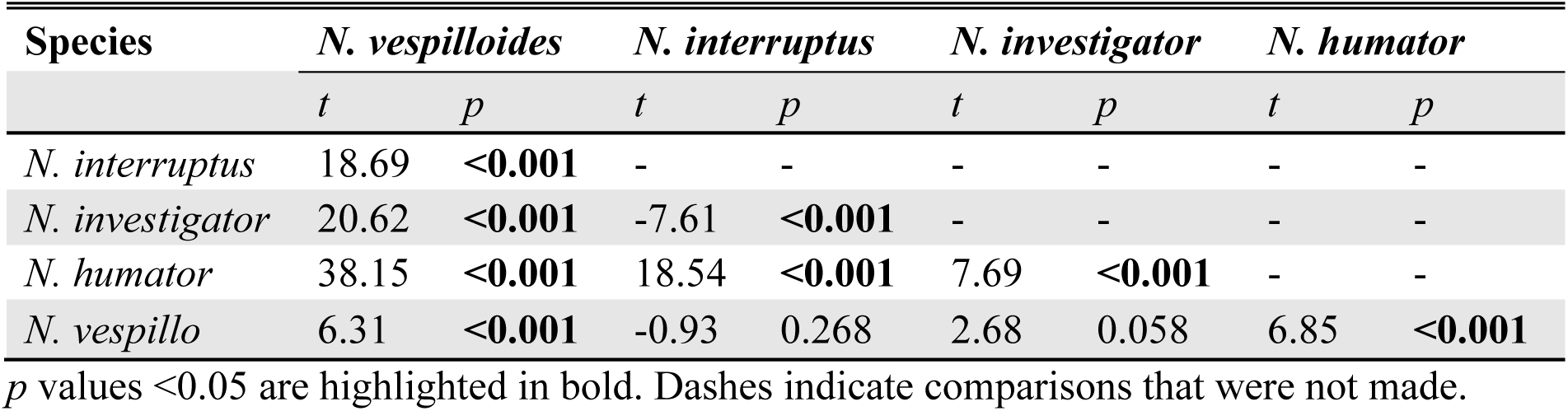
Post-hoc Tukey HSD comparing mean body size between *Nicrophorus* spp.

| Species | *N. vespilloides* |  | *N. interruptus* |  | *N. investigator* |  | *N. humator* |  |
| --- | --- | --- | --- | --- | --- | --- | --- | --- |
|  | *t* | *p* | *t* | *p* | *t* | *p* | *t* | *p* |
| *N. interruptus* | 18.69 | **<0.001** | - | - | - | - | - | - |
| *N. investigator* | 20.62 | **<0.001** | -7.61 | **<0.001** | - | - | - | - |
| *N. humator* | 38.15 | **<0.001** | 18.54 | **<0.001** | 7.69 | **<0.001** | - | - |
| *N. vespillo* | 6.31 | **<0.001** | -0.93 | 0.268 | 2.68 | 0.058 | 6.85 | **<0.001** |

*p* values <0.05 are highlighted in bold. Dashes indicate comparisons that were not made.

**Supplementary Table 3.** Results of the ANOVAs for division of carrion niche by *Nicrophorus* spp.

| **Dependent variable** | **Explanatory variables** | **$\chi^2$** | **d.f.** | ***p* value** |
| --- | --- | --- | --- | --- |
| Efficiency (%) | Carcass size | 17.26 | 1 | **<0.001** |
|  | Beetle species | 20.42 | 3 | **<0.001** |
|  | Carcass size x Beetle species | 28.85 | 3 | **<0.001** |

*p* values <0.05 are highlighted in bold.

**Supplementary Table 4.** Results of the ANOVAs for reaction norm experiment.

| **Dependent variable** | **Explanatory variables** | **$\chi^2$** | **d.f.** | ***p* value** |
| --- | --- | --- | --- | --- |
| Brood size | Carcass size | 3.33 | 1 | 0.068 |
|  | Population | 3.89 | 1 | **0.048** |
|  | Carcass size x Population | 4.80 | 1 | **0.029** |
| Average larval mass | Carcass size | 139.05 | 1 | **<0.001** |
|  | Population | 1.06 | 1 | 0.304 |
| Clutch size | Carcass size | 11.33 | 1 | **<0.001** |
|  | Population | 21.07 | 1 | **<0.001** |
|  | Female size | 4.26 | 1 | **0.039** |
| Average egg volume | Carcass size | 5.90 | 1 | **0.015** |
|  | Population | 4.59 | 1 | **0.032** |
| Total egg volume | Carcass size | 2.38 | 1 | 0.123 |
|  | Population | 2.16 | 1 | 0.141 |

*p* values <0.05 are highlighted in bold.

**Supplementary Table 5.** Differences in body size frequency distribution between *Nicrophorus* spp.

| Species | *N. vespilloides* |  | *N. interruptus* |  | *N. investigator* |  | *N. humator* |  |
| --- | --- | --- | --- | --- | --- | --- | --- | --- |
|  | *D* | *p* | *D* | *p* | *D* | *p* | *D* | *p* |
| *N. interruptus* | 0.54 | **<0.001** | - | - | - | - | - | - |
| *N. investigator* | 0.81 | **<0.001** | 0.41 | **<0.001** | - | - | - | - |
| *N. humator* | 0.91 | **<0.001** | 0.62 | **<0.001** | 0.38 | **<0.001** | - | - |
| *N. vespillo* | 0.70 | **<0.001** | 0.31 | 0.268 | 0.43 | 0.061 | 0.69 | **<0.001** |

D statistic and corresponding *p*-values from Kolmogorov-Smirnov tests. *p* values <0.05 are highlighted in bold. Dashes indicate comparisons that were not made.

**Supplementary Data 1. Spreadsheet file of gene set enrichment analysis results of the multiple GO terms for Gamlingay and Waresley *N. vespilloides*.**

## Notes

#### Summary of Updates

Several parts of the manuscript have been updated.

